# STIM1 at the plasma membrane: A novel Molecular regulator of B cell migration

**DOI:** 10.64898/2026.10.14.759286

**Authors:** Nelig Le Goux, Paul Buscaglia, Anthony Mainguy, Maela Hus, Mathieu Blery, Olivier Mignen

**Affiliations:** LBAI, UMR1227, Univ Brest, Inserm, Brest, France; Kalsiom SAS, 6 rue Porstrein, 29200 Brest, France; Antibody Engineering Platform, Queen Mary University of London Charterhouse Square, London, EC1M 6BQ; Department of Pharmacology, University of Michigan, Ann Arbor, MI, 48109, USA; Elizabeth Weiser Caswell Diabetes Institute, University of Michigan, Ann Arbor, MI, 48109, USA

## Abstract

Cell migration is a fundamental process required for effective immune responses, relying on both amoeboid movement and chemotaxis. Among chemotactic pathways, the CXCL12/CXCR4 axis plays a pivotal role in guiding B cell migration. While essential under physiological conditions, this pathway is frequently dysregulated in pathological contexts such as Chronic Lymphocytic Leukemia (CLL) and Systemic Lupus Erythematosus (SLE), where altered CXCR4 expression enhances B cell retention, survival, and tissue infiltration. Calcium signaling is central to B cell migration, notably through Store-Operated Calcium Entry (SOCE), STIM1 located at the endoplasmic reticulum (ER) membrane and Orai1 calcium channels. Although STIM1 is traditionally studied as an ER protein regulating SOCE, early findings and recent work from our group indicate that STIM1 is also present at the plasma membrane (STIM1_PM_), where it displays a dual topology and contributes to constitutive calcium entry (CCE). The function of STIM1_PM_ pool in B cell migration remains unknown. Here, we demonstrate that both STIM1_ER_ and Orai1 are required for CXCL12-induced migration of B cells. Importantly, we provide the first evidence that STIM1_PM_, exhibiting N-terminal and C-terminal orientations, actively regulates B cell migration independently of SOCE. We show that STIM1_PM_ controls CCE and modulates intercellular interactions, thereby influencing chemotactic responses. Targeting STIM1_PM_ domains significantly impairs B cell migration, highlighting its potential as a novel therapeutic target in diseases characterized by aberrant B cell trafficking. Our findings uncover an unanticipated role for STIM1PM and open new perspectives on calcium-dependent and calcium-independent mechanisms of B cell migration.

## Introduction

Cell migration is an essential physiological phenomenon for an effective adaptive immune response. Two types of cell migration can be described: amoeboid migration and chemotaxis. Amoeboid migration characterizes the rapid movement of isolated cells. Various cell types are capable of performing amoeboid migration, including B cells, T cells, dendritic cells, and neutrophils (Lammermann and Sixt, 2009). Amoeboid cells use friction forces to transform their internal movements into migration (Lammermann and Sixt, 2009). Chemotaxis is defined as the induction of migration in response to a soluble phase chemokine concentration gradient. This migration is oriented from the low concentration area to the higher concentration area (Worbs et al., 2017). Additionally, transendothelial migration of B cells allow their migrate through endothelial cells to move from the bloodstream to secondary lymphoid organs (SLOs).

Chemokines are a family of small soluble proteins ranging from 8 to 14 kDa, divided into four subfamilies: CCL, CXCL, CX3CL, and XCL (Nomiyama et al., 2013) that bind to specific receptors, which are also divided into four subfamilies: CCR, CXCR, CX3CR, and XCR (Hughes and Nibbs, 2018). Chemokines are majors’ actors of leucocyte cell migration (Hughes and Nibbs, 2018). Our study focuses on CXCL12 chemokine capable of inducing B cell migration through chemotaxis. CXCL12 is the only ligand for CXCR4, its receptor expressed on the surface of various cell types such as lymphocytes in the bone marrow, and in lymphoid organs but also in kidneys and liver (Mousavi, 2020). It is known for a long time that CXCL12 is constitutively produced by stromal cells in the bone marrow, allowing the retention of hematopoietic progenitors in the bone marrow (Peled et al., 1999). Additionally, CXCL12 plays a major role in the transmigration of B cells through high endothelial venules (HEV) (Phillips and Ager, 2002) or through cell lines (HUVEC) (Cinamon et al., 2001; Grabovsky et al., 2000). While CXCR4 and CXCL12 are essential under physiological conditions, numerous studies have focused on their deregulation in pathological conditions, particularly autoimmune diseases and cancers (Marchese and Benovic, 2001; Werner et al., 2013). In these pathologies, cell migration, especially B cell migration, can be deregulated.

In Chronic Lymphocytic Leukemia (CLL), characterized by the accumulation of monoclonal CD5⁺ B cells in the blood, bone marrow, and secondary lymphoid organs (Swerdlow et al., 2016), B cells exhibit impaired migratory capacity compared to those from healthy individuals. Pathophysiological processes alter B cell properties such that, while retaining their ability to recognize chemokine receptors, they exploit the overexpression of these receptors to enhance migration toward secondary lymphoid organs, leading to the accumulation of malignant B cells in these tissues (Holroyd and Michie, 2018). For instance, CXCR4 is overexpressed in CLL B cells (Burger et al., 1999), as are other chemokine receptors such as CXCR5, which binds to CXCL13 (Burkle et al., 2007).

In systemic lupus erythematosus (SLE), B cells are hyperactivated and differentiate into autoantibody-secreting plasma cells (Chong and Mohan, 2009). Similar to CLL, B cell migration defects are observed in association with the CXCR4/CXCL12 axis. In lupus-prone mouse models, CXCR4 is overexpressed on B cells, enhancing their migration and survival. Elevated CXCR4 expression has also been reported in B cells from SLE patients (Wang et al., 2009). Moreover, CXCR4 overexpression in SLE B cells positively correlates with disease activity (Zhao et al., 2017). As in CLL, this overexpression promotes increased migration to secondary lymphoid organs and enhanced survival, thereby facilitating the differentiation of autoreactive B cells into autoantibody-secreting plasma cells. Consequently, several studies have targeted the CXCR4/CXCL12 pathway in murine models of SLE as a potential therapeutic strategy (Chong and Mohan, 2009).

The calcium ion is a crucial element for cellular physiology, particularly involved in cell migration, (Hammad et al., 2021, DerMardirossian et al., 2024). The role of STIM1 (Stromal Interaction Molecule 1) in cell migration is well known and primarily described through the prism of the ubiquitous and well described Store-Operated Calcium Entry (SOCE). STIM1 and SOCE are described as having pro-migratory and pro-invasive activities in various cancers, including breast cancer (Feng et al., 2010), hepatocellular carcinoma (Yang et al., 2013), nasopharyngeal carcinoma cell (Luo et al., 2023, Deng et al., 2025) colorectal cancer (Wong and Chang, 2015) and melanoma (Fedida-Metula et al., 2012).

Cell migration induced by CXCL12/CXCR4 is calcium-dependent and involved STIM1 and Orai1 protein. The binding of CXCL12 to its receptor CXCR4 causes an increase in intracellular calcium concentration in various cell types (Janssens et al., 2016). In B cells, the activation of CXCR4 by CXCL12 triggers the release of endoplasmic reticulum calcium stores and the subsequent SOCE, essential for B cell migration (Latour et al., 2018). The activation of CXCR4 induces the cleavage of PIP2 (phosphatidylinositol (3,4)-bis-phosphate) into InsP3 (inositol trisphosphate) and DAG (diacylglycerol) through the activation of PLCγ2 (phospholipase C gamma 2). Activation of InsP3 receptors (InsP3R) on the endoplasmic reticulum (ER) membrane, cause a release of the cell’s calcium -stores sensed by STIM1 located on the ER membrane. STIM1 activation when ER Ca^2+^ is lowered leads to its, oligomerization, conformational change and translocation to ER/PM contact sites allowing its interaction with ORAI calcium channels on the plasma membrane responsible for the calcium influx from the extracellular environment (SOCE) (Feske et al., 2012, Rashwan et al., 2021, <u>Kodakandla</u> et al., 2023)

STIM1 is described as a protein primarily located on the ER membrane, and its involvement in cell migration is only considered in this context and its role in regulating and activating SOCE influx. However, early studies mentioning STIM1 describe this protein as being located with a type I orientation on the plasma membrane (PM) of pre-B cells and involved in the interaction with stromal cells (Manji et al., 2000; Oritani and Kincade, 1996; Williams et al., 2001). These initial studies attributed a role for STIM1 located in PM (STIM1_PM_) in cell-cell interactions. Since then, it has been established that STIM1_PM_ is also involved in regulating calcium influxes: ARC (Mignen et al., 2007) and constitutive calcium entry (ECC) (Debant et al., 2019). Contrary to initial suggestions (Jardin et al., 2009; Rao et al., 2010; Spassova et al., 2006), it seems that STIM1_PM_ does not regulate SOCE-type influx (Debant et al., 2019). In addition, recent studies conducted by our team have shown that STIM1_PM_ is overexpressed in B cells from CLL patients (Debant et al., 2019). Furthermore, STIM1_PM_ is inserted into the plasma membrane of cells with a dual topology, with both the N-terminal domain (N-Ter out, type I topology) or the C-terminal domain (C-Ter out type II topology) exposed to the extracellular environment (Buscaglia et al., 2026).

As previously mentioned, Latour’s team highlighted the role of STIM1 in basal or CXCL12 induced DLBCL cell migration. Although STIM1 and Orai1 are responsible for Ca^2+^ influx induced by CXCL12, cell migration induced by this chemokine is independent of Ca^2+^ entry. This Ca^2+^-independent but STIM1-dependent signaling pathway involved in basal and CXCR4 dependent cell migration, may be linked to the coupling of RhoA and ROCK dependent signaling pathway to MLC2 phosphorylation and actin polymerization (Latour 2018). Only the fraction of STIM1 located in the ER membrane is considered in this study. Based on known role of STIM1 it is still difficult to explain the exact role of STIM1 in this Ca^2+^-independent migration mechanism. We hypothesize therefore hat STIM1_PM_ is involved in cell migration through its role in regulating calcium influx independent of intracellular calcium store release.

The involvement of STIM1 in cell migration and the increased expression of STIM1_PM_ in cancer B cells could be associated to the migration deregulation observed in these cells. It is therefore important to determine the role of STIM1_PM_ in B cell migration induced by CXCL12 to highlight the potential therapeutic role of STIM1_PM_.

The results obtained in our study confirm that STIM1_RE_, Orai1, and by extension SOCE influx are important actors in B cell migration induced by CXCL12. For the first time, it is demonstrated that STIM1_PM_ presents a dual N-Terout and C-Terout topology on the plasma membrane of B cells and that STIM1_PM_ regulates the migration of these cells in vitro. The results confirm that STIM1_PM_ is involved in regulating ECC without participating in SOCE-type influxes. The inhibition of B cell migration by molecules targeting the N and C domains of the protein confirms the therapeutic perspectives offered by biotherapies targeting STIM1_PM_. Our work suggests that STIM1_PM_ modulates cell migration through its action on calcium influxes and intercellular interactions.

## Results

### SOCE Influx Regulates CXCL12-Induced Migration of JOK-1 B Cells

Lymphocyte migration is a complex process involving multiple molecular actors, including chemokines (Hughes and Nibbs, 2018). CXCL12 and its receptor CXCR4 are key regulators of B cell migration under both physiological and pathological conditions (Cojoc et al., 2013 Zhou et al., 2023, Korbecki et al., 2024), acting through intracellular signaling cascades such as the PLC, MAPK and PI3K/CDC42 pathways, as well as cytoskeletal remodeling (Chatterjee et al., 2014). SOCE influx mediated by STIM1 is considered essential for B cell migration in response to a CXCL12 gradient.

The B cell lines JOK-1 and RAMOS, which express CXCR4 on their plasma membrane, were selected for migration assays (Supplementary Figure 1A and 1B). In a transwell migration model mimicking chemokine gradient-induced migration, approximately 30% of JOK-1 B cells migrated in response to CXCL12 (200 ng/mL) (Figure 1A). Similarly, trans-endothelial migration of JOK-1 cells across a monolayer of human umbilical vein endothelial cells (HUVECs) reached approximately 30% (Figure 1A). We also confirmed that CXCR4 undergoes internalization upon CXCL12 binding in JOK-1 cells (Supplementary Figure 1C).

**Figure 1.**
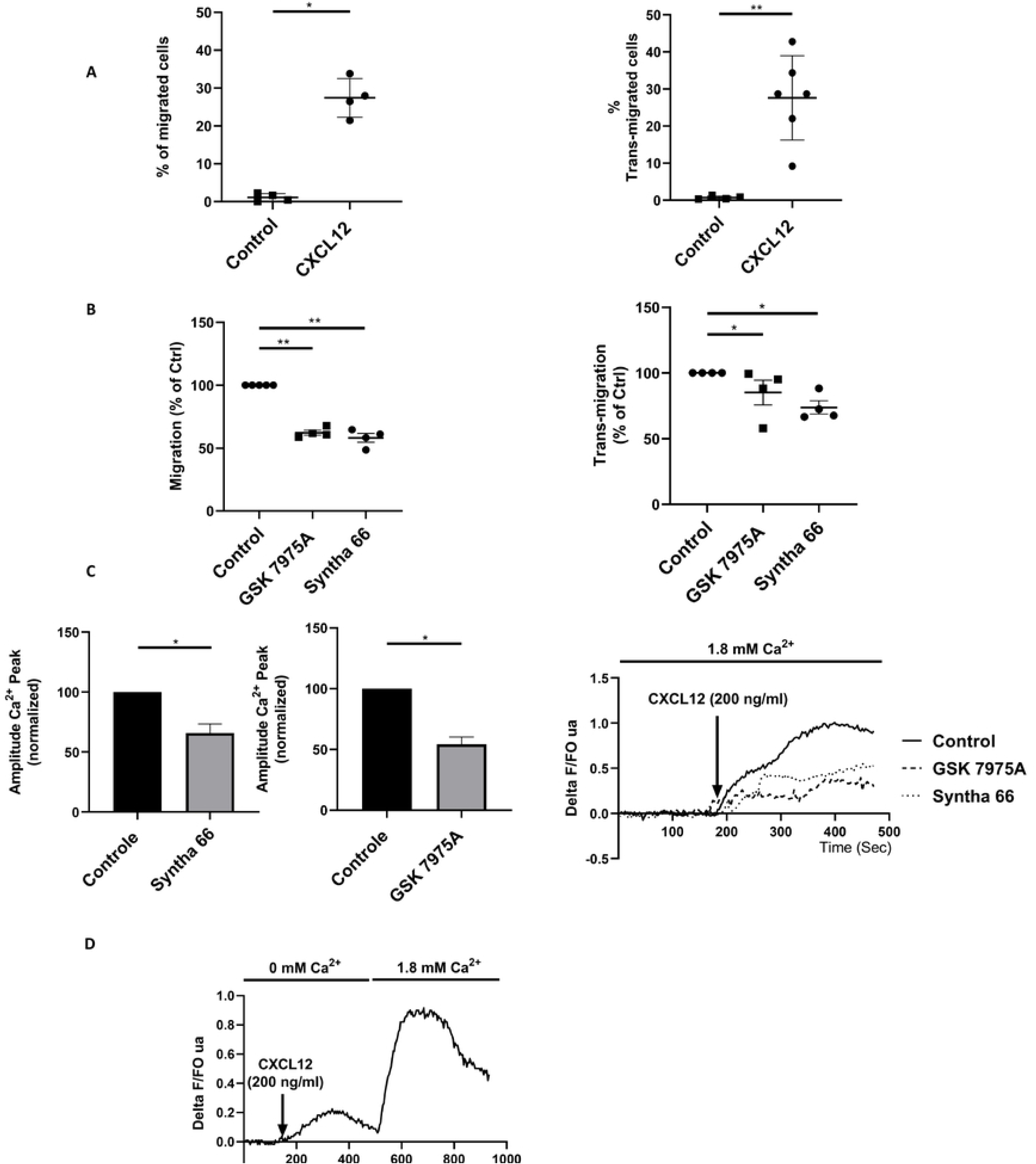
CXCL12-Induced Migration of JOK-1 Cells Is Regulated by Orai1. **A.** Percentage of JOK-1 cell migration in response to a CXCL12 gradient (200 ng/mL) through a transwell filter (left panel, *N*=4) and across an endothelial monolayer (right panel, *N*=6). Histograms represent the percentage of migrated cells normalized to control conditions (no chemokine gradient). **B.** Effect of Orai1 inhibitors (GSK 7975A, 5 µM, *N*=4; Syntha 66, 5 µM, *N*=4) on CXCL12-induced migration (200 ng/mL in the lower chamber) of JOK-1 cells through a transwell filter (left panel) or across HUVEC endothelial cells (right panel). Migration percentages are normalized to control conditions (CXCL12 + 0.1% DMSO). **C.** Effect of Orai1 inhibitors (GSK 7975A, 5 µM; Syntha 66, 5 µM) on the amplitude of the calcium signal induced by CXCL12 (200 ng/mL) in Fura-2-loaded JOK-1 cells. Histograms (left panel) show normalized calcium signal amplitudes relative to vehicle-treated controls (0.1% DMSO). Representative calcium traces (right panel) illustrate responses in the presence or absence of inhibitors. **D.** Representative trace of the calcium signal induced by CXCL12 (200 ng/mL), recorded first in calcium-free medium and then after the addition of 1.8 mM extracellular Ca²⁺. Data were analyzed using a non-parametric *t*-test. *P*<0.05 (*), P<0.01 (**), P*<0.001 (\*\*\**).

As shown in Figure 1B, pharmacological inhibition of Orai1 using specific inhibitors (GSK 7975A and Syntha 66) significantly reduced both transwell migration (GSK 7975A: 38% inhibition, *p*<0.001; Syntha 66: 42% inhibition, *p*<0.001) and trans-endothelial migration (GSK 7975A: 15% inhibition, *p*<0.005; Syntha 66: 25% inhibition, *p*<0.005) of JOK-1 B cells in response to a CXCL12 gradient. These latter results suggest the involvement of the calcium channel Orai1 in efficient cell migration. This impaired migration is likely due to the inhibition of the CXCL12-induced calcium signal, as evidenced by a significant reduction in calcium signal amplitude (GSK 7975A: 35% inhibition, *p*<0.005; Syntha 66: 45% inhibition, *p*<0.005) in JOK-1 B cells treated with these Orai1 inhibitors (Figure 1C). As previously described in Latour’s work (Latour et al., 2018), CXCL12 induces an ORAI1-dependent calcium influx (Figure 1C), but contrary to the observations made in their study, the migration of JOK1 B cells also appears to depend on ORAI1 (Figure 1B). Altogether, our data reveal that unlike previous observations in other B cell models, JOK-1 cells rely on Orai1-dependent calcium influx for CXCL12-induced migration, underscoring cell-type-specific differences in calcium signaling requirements.

### STIM1 Is Present at the Plasma Membrane of JOK-1 B Cells in Two Distinct Topologies

Our work, along with that of other groups, has reported the presence of STIM1_PM_ exhibiting a type I membrane topology, characterized by an extracellularly oriented N-terminus (N-Ter_out_) (Debant et al., 2019; Manji et al., 2000; Mignen et al., 2007; Oritani and Kincade, 1996; Williams et al., 2001). More recently, we demonstrated that STIM1_PM_ can adopt a dual topology (both N-Ter_out_ and C-Ter_out_) when present at the cell surface (Buscaglia et al., 2026).

Therefore, we evaluate the presence and dual topology of STIM1_PM_ in B lymphocytes and endothelial cells for understanding the role of this fraction of STIM1 in CXCL12-induced B cell migration. Initially, the presence of STIM1_PM_ with an N-Ter_out_ orientation at the plasma membrane of JOK-1 cells was detected by flow cytometry using monoclonal antibodies targeting the N-terminal domain of STIM1 (N-Ter 1: clone GOK, BD; N-Ter 2: clone BY-12; N-Ter 3: clone BM-4, Diaclone) (Figure 2A), with epitope locations shown in Supplementary Figure 2B. Similar results were obtained with the RAMOS B cell line (Supplementary Figure 2), although the lower expression of N-Ter_out_ STIM1_PM_ in these cells led to their exclusion from subsequent experiments.

**Figure 2.**
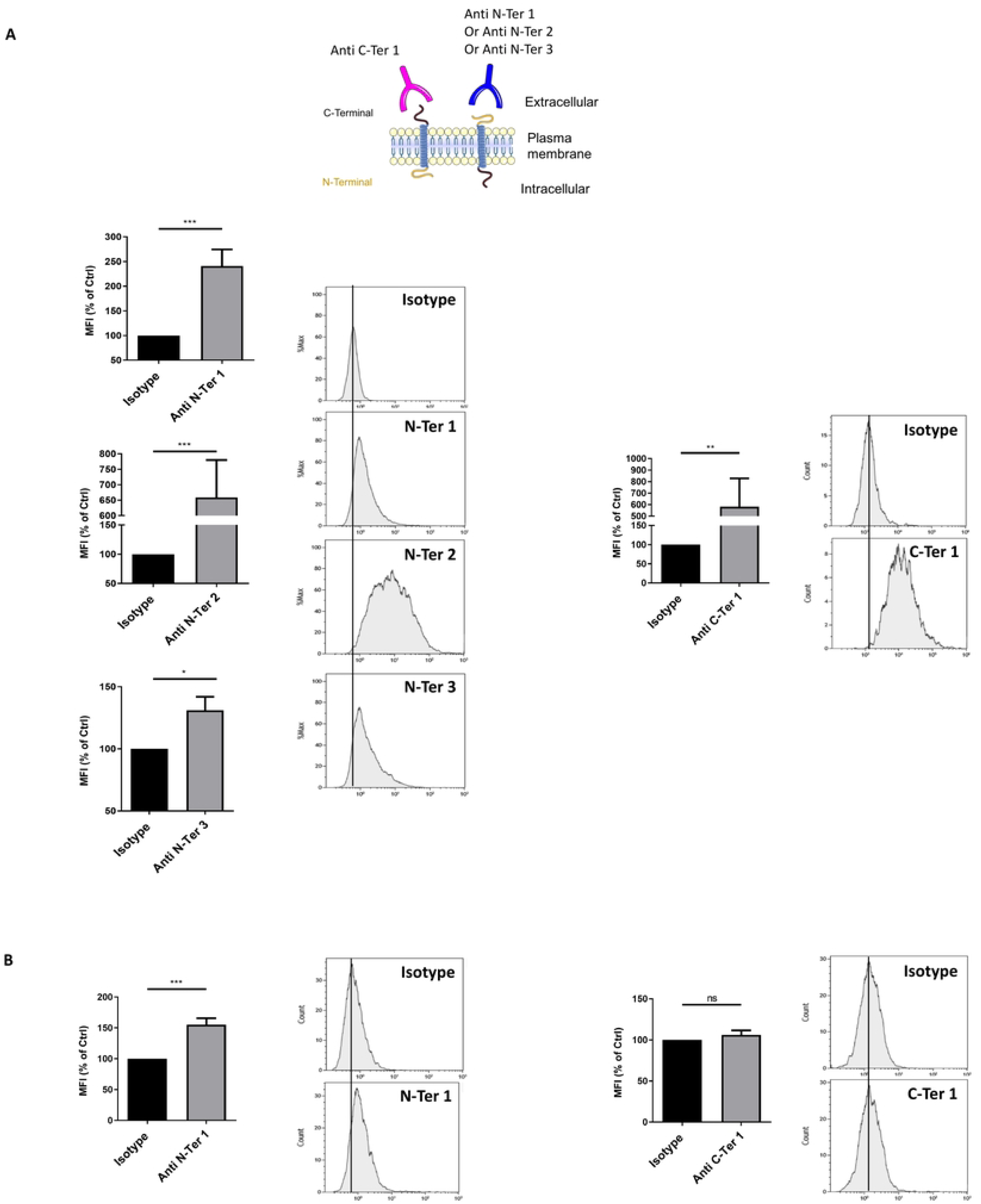
STIM1_PM_ Is Present at the Plasma Membrane of JOK-1 Cells and Exhibits Dual Topology. **A–B.** The presence and membrane topology of STIM1_PM_ were assessed by flow cytometry in non-fixed, non-permeabilized JOK-1 cells (**Panel B**) and HUVEC endothelial cells (**Panel C**). Cells were incubated with monoclonal anti-STIM1 antibodies targeting either the N-terminal domain (left panels: anti-STIM1 N-Ter 1, clone GOK-PE, BD; N-Ter 2, clone BY-12-PE, Diaclone; N-Ter 3, clone BM-4-PE, Diaclone) or the C-terminal domain (right panels: anti-STIM1 C-Ter 1, clone CDN3H4-PE, Santa Cruz). For each experiment (*N*>4), the median fluorescence intensity (MFI ± SEM) was normalized to the MFI of cells incubated with the corresponding isotype control. Representative overlay histograms are shown for each antibody condition. Statistical analysis was performed using a non-parametric *t*-test. *P*<0.05 (*), P<0.01 (**), P*<0.001 (\*\*\**).

Importantly, for the first time in B cell lines (JOK-1), we observed STIM1_PM_ with a C-Ter_out_ orientation. In contrast, RAMOS B cells exhibited only low levels of C-Ter_out_ STIM1_PM_, further supporting their exclusion from further analyses (Figure 2A and Supplementary Figure 2). In HUVEC endothelial cells, STIM1_PM_ was also detected, but exclusively in a type I topology, as suggested by flow cytometry data (Figure 2B). The percentage of positive cells and raw mean fluorescence intensity (MFI) values for each antibody and cell line are summarized in Table 1. These data reinforce the decision to focus on JOK-1 B cells, which exhibit stronger STIM1_PM_ amount.

| Cell Lines | MFI Brute moyenne |  |  |  | % of positives cells |  |  |  |
| --- | --- | --- | --- | --- | --- | --- | --- | --- |
|  | Anti N-Ter 1 | Anti N-Ter 2 | Anti N-Ter 3 | Anti C-Ter 1 | Anti N-Ter 1 | Anti N-Ter 2 | Anti N-Ter 3 | Anti C-Ter 1 |
| JOK-1 | 2,4 ± 0,7<br>N=8 | 6,5 ± 1,2<br>N=8 | 1,3 ± 0,1<br>N=8 | 5,8 ± 2,6<br>N=8 | 85,5% | 96,90% | 78,5% | 98,72% |
| RAMOS | 1,1 ± 0,1<br>N=7 | 4,6 ± 0,9<br>N=7 | 1,3 ± 0,1<br>N=7 | 1,2 ± 0,2<br>N=7 | 4,50% | 90,20% | 65,90% | 31,80% |
| HUVEC | 1,5 ± 0,2<br>N=5 |  |  | 1,1 ± 0,1<br>N=5 | 59,10% |  |  | 13,68% |

Thus, the STIM1_PM_ fraction in the two B cell lines studied displays a dual topology (N-Ter_out_ and C-Ter_out_). This dual topology will be taken into account in the subsequent investigation of STIM1_PM_ involvement in CXCL12-dependent B cell migration.

### Constitutive Calcium Entry in JOK-1 B Cells Is Regulated by Plasma Membrane-Associated STIM1 Regardless of Its Topology

While the role of ER-resident STIM1 (STIM1_ER_) in calcium influx, particularly in store-operated calcium entry (SOCE), is well established and extensively documented (Dai et al., 2024, Ambudkar et al., 2017, Collins et al., 2013), the involvement of STIM1_PM_ in calcium signaling remains poorly explored. However, it has been shown that blocking STIM1_PM_ using an antibody targeting its N-terminal domain inhibits calcium influxes that are independent of ER store depletion, such as arachidonate-regulated calcium (ARC) entry and constitutive calcium entry (CCE), without affecting SOCE (Mignen et al., 2007; Debant et al., 2019).

In JOK-1 B cells, application of a blocking antibody against the N-terminal domain of STIM1_PM_ significantly reduced CCE by 12% (*p*<0.001, *N*=6), confirming the involvement of STIM1_PM_ in this process. Similarly, an antibody targeting the C-terminal domain of STIM1 also significantly inhibited CCE by 20% (*p*<0.001, *N*=5) (Figure 3A). These results indicate that STIM1_PM_ regulates CCE in these cells regardless of its membrane orientation (N-Ter_out_ or C-Ter_out_). Furthermore, blocking STIM1_PM_ with either N- or C-terminal antibodies did not affect ER calcium release (Supplementary Figure 3A) and SOCE amplitude (Figure 3B). Consistent with previous findings (Debant et al., 2019), Orai1 also also contributes to CCE, as specific inhibitors (GSK 7975A and Syntha 66) significantly reduced both CCE amplitude (Figure 3C) and SOCE (Supplementary Figure 3B) in JOK-1 B cells. We also hypothesize that other calcium channels, such as TRPC1, may be involved in the regulation of constitutive calcium entry.

**Figure 3.**
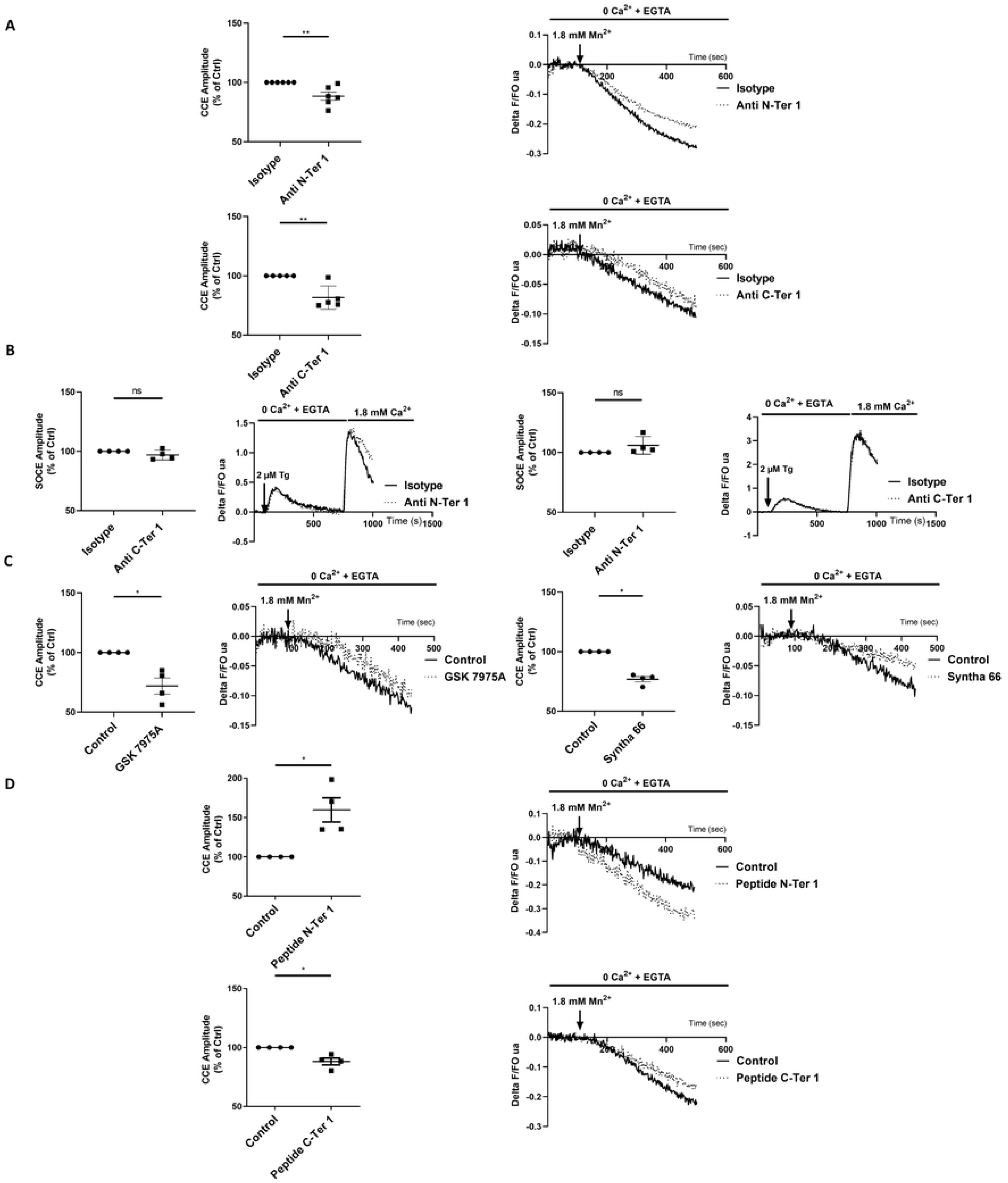
STIM1_PM_ and Orai1 Modulate Constitutive Calcium Entry in JOK-1 Cells. Constitutive calcium entry (CCE) was estimated by measuring the rate of Fura-2 fluorescence quenching by extracellular Mn²⁺ in calcium-free medium. SOCE was induced by ER calcium store depletion using thapsigargin (2 µM), followed by the addition of 1.8 mM extracellular Ca²⁺. **A.** Effect of anti-STIM1 antibodies targeting the N-terminal domain (top panel: anti-STIM1 N-Ter 1, clone GOK, BD, 10 µg/mL; *N*=6) or the C-terminal domain (bottom panel: anti-STIM1 C-Ter 1, clone CDN3H4, Santa Cruz, 10 µg/mL; *N*=5) on CCE amplitude in JOK-1 cells. **B.** Effect of anti-STIM1 N-Ter 1 (left panel) and C-Ter 1 (right panel) antibodies (10 µg/mL; *N*=4 each) on SOCE amplitude in JOK-1 cells. **C.** Quantification of the effect of Orai1 inhibitors on CCE amplitude: GSK 7975A (5 µM, *N*=4, left panel) and Syntha 66 (5 µM, *N*=4, right panel). **D.** Effect of STIM1-derived peptides on CCE amplitude: N-Ter 1 peptide (aa 63–200, 10 µg/mL, *N*=4, top panel) and C-Ter 1 peptide (aa 637–672, 10 µg/mL, *N*=4, bottom panel). Cells were incubated with antibodies, peptides, or inhibitors for 1 hour. Amplitude values (individual data points ± SEM) were normalized to control conditions (isotype control, vehicle, or no peptide). Representative traces of calcium signals for each condition are shown. Statistical analysis was performed using a non-parametric *t*-test. *P*<0.05 (*), P<0.01 (**), P*<0.001 (\*\*\**).

Finally, the effects of synthetic peptides corresponding to specific STIM1 domains were tested on both SOCE and CCE. The peptides used are detailed in Supplementary Figure 2B. Interestingly, a peptide derived from the N-terminal domain of STIM1 (amino acids 63–200) significantly increased CCE amplitude by 60% (*p*<0.005, *N*=4), whereas a peptide corresponding to amino acids 637–672 of the C-terminal domain of the protein reduced CCE by 15% (*p*<0.005, *N*=4) (Figure 3D). As observed with the antibodies, these peptides had no significant effect on ER calcium release and SOCE amplitude (Supplementary Figure 3C).

Taken together, these results confirm that N-Ter_out_-oriented STIM1_PM_ contributes to CCE. Moreover, for the first time, we demonstrate that STIM1_PM_ with a type II topology (C-Ter_out_) also regulates CCE in B cells. However, STIM1_PM_, regardless of its membrane orientation, does not appear to participate in the regulation of SOCE.

### STIM1_PM_ Modulates CXCL12-Induced Migration of JOK-1 B Cells Without Affecting the Specific Chemokine Associated Calcium Signal

As mentioned, Stimulation of CXCR4 by CXCL12 triggers ER calcium release via InsP3 receptor activation, subsequent activation of ER-resident STIM1 (STIM1_ER_), opening of Orai1 channels, and a robust store-operated calcium entry (SOCE) from the extracellular space (Latour et al., 2018; Zuccolo et al., 2018). We observed such a SOCE influx in JOK-1 cells upon CXCL12 stimulation (Figure 1C). It is proposed that CXCL12/CXCR4-induced cell migration is regulated by calcium signaling involving STIM1 and Orai1 and suggested that STIM1 contributes to calcium independent regulation of this type of migration (WenChen et al., 2016, Maltan et al., 2023).

In our study, Orai1 inhibitors suppressed both SOCE and the calcium signal triggered by CXCL12 in JOK-1 cells (Figure 1C and Supplementary Figure 3B). In contrast, treatment with anti-STIM1 antibodies or STIM1-derived peptides did not affect SOCE (Figure 3B and Supplementary Figure 3C) but did modulate constitutive calcium entry (CCE) (Figures 3A and 3D). Surprisingly, the N-terminal peptide increases constitutive calcium entry from 50%, whereas the C-terminal peptide decreases CCE from 15%. Moreover, neither N-terminal nor C-terminal STIM1-targeting antibodies or peptides altered the calcium signal induced by CXCR4 stimulation (Figure 4A). Collectively, these results confirm that STIM1_PM_ is not involved in SOCE and that CXCL12-induced SOCE is independent of STIM1_PM_ regulation.

**Figure 4.**
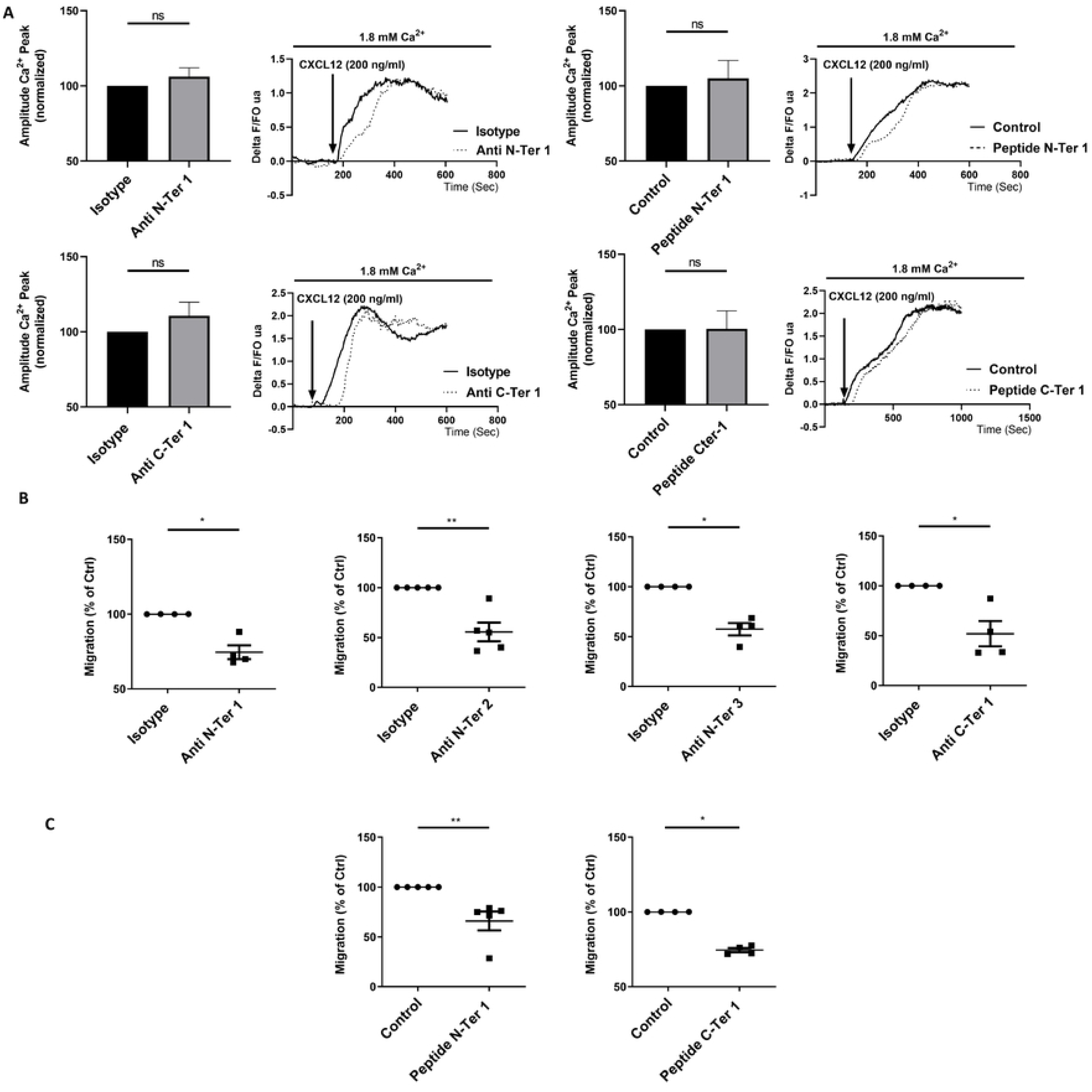
Anti-STIM1 Antibodies and STIM1 Peptides Inhibit CXCL12-Induced Migration of JOK-1 Cells Without Affecting the Associated Calcium Signal. **A.** Effect of anti-STIM1 antibodies (top panel: anti-STIM1 N-Ter 1, clone GOK, BD, 10 µg/mL, *N*=4; anti-STIM1 C-Ter 1, clone CDN3H4, Santa Cruz, 10 µg/mL, *N*=4) and STIM1-derived peptides (bottom panel: N-Ter 1, aa 63–200, 10 µg/mL, *N*=4; C-Ter 1, aa 637–672, 10 µg/mL, *N*=4) on the calcium signal induced by CXCL12 (200 ng/mL) in Fura-2-loaded JOK-1 cells. Cells were incubated for 1 hour with or without antibodies or peptides. Histograms show normalized calcium signal amplitudes relative to control conditions (isotype control or no peptide). Representative calcium traces are shown for each condition. **B–C.** Effect of anti-STIM1 antibodies (**Panel B**) and STIM1 peptides (**Panel C**) on JOK-1 cell migration through a transwell filter in response to a CXCL12 gradient (200 ng/mL, 12 h). Antibodies: anti-STIM1 N-Ter 1 (clone GOK, BD), N-Ter 2 (clone BY-12, Diaclone), N-Ter 3 (clone BM-4, Diaclone), and C-Ter 1 (clone CDN3H4, Santa Cruz), all at 10 µg/mL. Peptides: N-Ter 1 (aa 63–200) and C-Ter 1 (aa 637–672), both at 10 µg/mL. Migration percentages are normalized to control conditions (isotype or no peptide). Statistical analysis: non-parametric *t*-test. *P*<0.05 (*), P<0.01 (**), P*<0.001 (\*\*\**).

Given that STIM1_PM_ regulates Orai1-mediated CCE and that Orai1 inhibition significantly reduces CXCL12-induced chemotactic migration of JOK-1 B cells (Figure 1B), we further investigated the role of STIM1_PM_ in this process. First, CXCL12 treatment did not alter STIM1_PM_ expression levels in JOK-1 cells (Supplementary Figure 4A). However, all three N-terminal STIM1-targeting antibodies significantly inhibited JOK-1 migration through a transwell filter by 25% (anti-N-Ter 1), 45% (anti-N-Ter 2), and 40% (anti-N-Ter 3), respectively (Figure 4B). Similarly, the C-terminal STIM1 antibody (anti-C-Ter 1) reduced migration by approximately 50% (Figure 4B). In addition, STIM1-derived peptides targeting the N-terminal (N-Ter 1) and C-terminal (C-Ter 1) domains also significantly inhibited migration by 35% and 25%, respectively (Figure 4C). These peptide effects are unexpected, given that the N-terminal peptide enhances CCE while the C-terminal peptide reduces it. The reason why similar effects on cell migration are not reflected in ECC remains unclear.

Altogether, these findings reveal that STIM1_PM_ modulates CXCL12-induced migration of JOK-1 B cells through mechanisms independent of its role in SOCE or CXCR4-triggered calcium signaling, highlighting a distinct regulatory pathway that remains to be fully elucidated.

### Trans-Endothelial Migration of JOK-1 B Cells Is Also Regulated by Plasma Membrane-Associated STIM1

As previously shown, CXCL12 gradient -induced migration of JOK-1 cells through a transwell filter is inhibited by blocking STIM1_PM_, regardless of its membrane orientation. We next focused on the trans-endothelial migration of JOK-1 cells across a monolayer of HUVECs in response to a CXCL12 gradient. Our results demonstrate that targeting the N-terminal domain of STIM1_PM_ located on B cells (preincubation of JOK-1 only) either with anti-STIM1 antibodies or with N-terminal STIM1-derived peptides-specifically is sufficient to significantly inhibit their trans-endothelial migration (Figure 5A). The degree of inhibition varied among the antibodies: anti-STIM1 N-Ter 1 reduced migration by 16%, N-Ter 2 by 20%, and N-Ter 3 by 30% (Figure 5A). A dose-dependent inhibition was observed when cells are incubated with the N-Ter 1 peptide, with effects starting at 0.5 µg/mL and reaching a maximum of approximately 40% inhibition at 10 µg/mL (Figure 5B).

**Figure 5.**
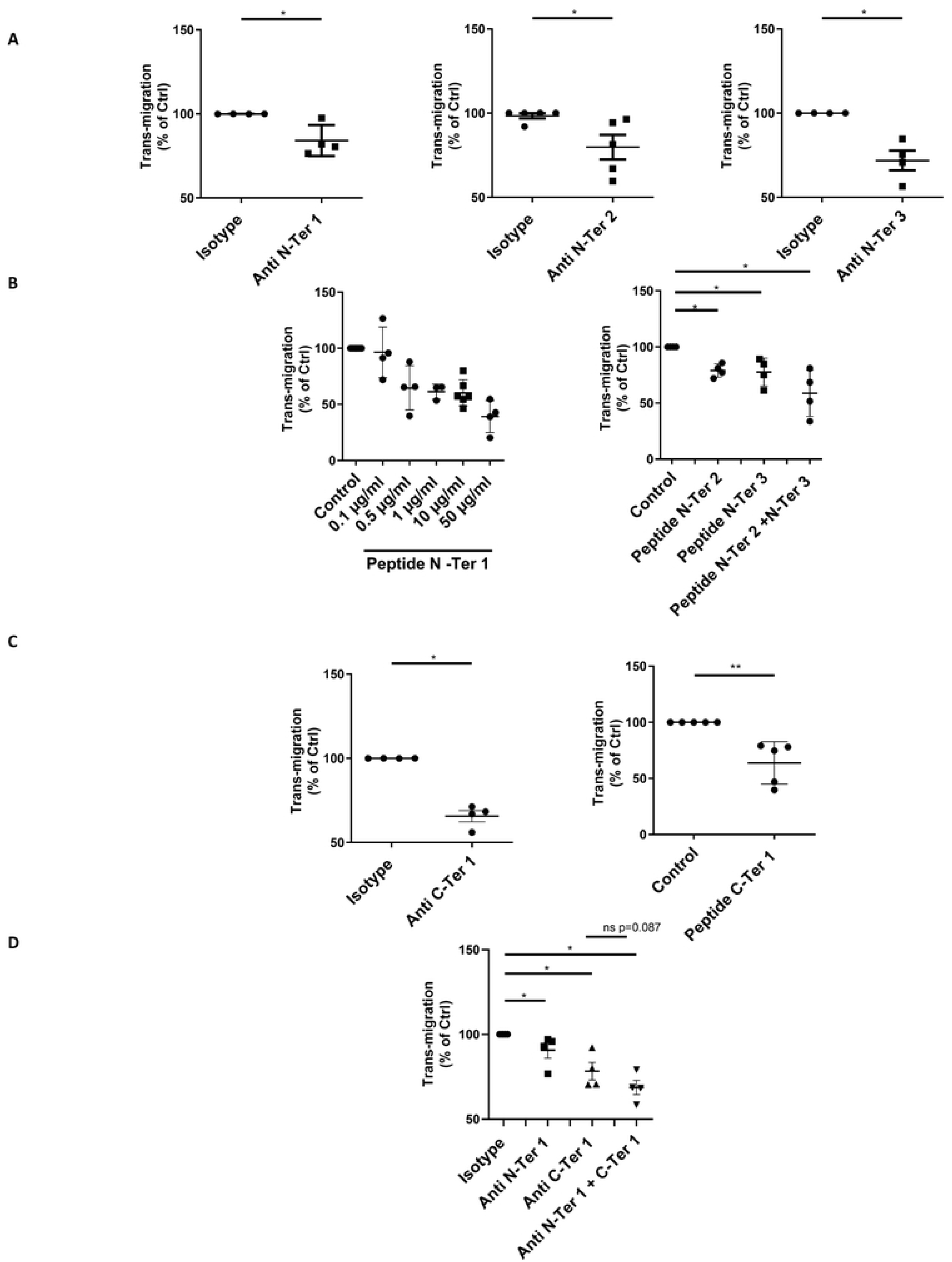
Anti-STIM1 Antibodies and STIM1 Peptides Inhibit CXCL12-Induced Trans-Endothelial Migration of JOK-1 Cells. For all migration assays, B cells were pre-incubated for 1 hour with antibodies or peptides. **A.** Effect of anti-STIM1 antibodies targeting the N-terminal domain: N-Ter 1 (clone GOK, BD), N-Ter 2 (clone BY-12, Diaclone), and N-Ter 3 (clone BM-4, Diaclone), all at 10 µg/mL (*N*=4–5). **B.** Effect of STIM1 N-terminal peptides: N-Ter 1 (aa 63–200, 0.1–50 µg/mL, *N*=4–8), N-Ter 2 (aa 128–168, 10 µg/mL, *N*=4), and N-Ter 3 (aa 96–127, 10 µg/mL, *N*=4). **C.** Effect of anti-STIM1 C-Ter 1 antibody (clone CDN3H4, Santa Cruz, 10 µg/mL, *N*=4) and C-Ter 1 peptide (aa 637–672, 10 µg/mL, *N*=4) on trans-endothelial migration. **D.** Additive effect of combined anti-STIM1 N-Ter 1 (2.5 µg/mL) and C-Ter 1 (2.5 µg/mL) antibodies on JOK-1 trans-endothelial migration. Cells were pre-incubated for 12 hours before contact with HUVECs (*N*=4 per condition). Data are presented as normalized migration percentages (individual values ± SEM) relative to control conditions (isotype or no peptide). Statistical analysis: non-parametric *t*-test. *P*<0.05 (*), P<0.01 (**), P*<0.001 (\*\*\**).

Binding of the N-Ter 1 peptide to membrane-associated STIM1 in JOK-1 cells was confirmed by flow cytometry (Supplementary Figure 4B). The inhibitory effect of N-terminal peptides on trans-endothelial migration varied depending on their sequence. Peptides N-Ter 2 (aa 128–168) and N-Ter 3 (aa 96–127), corresponding to the EF-hand and SAM domains respectively, are both contained within the broader N-Ter 1 fragment (aa 63–200). Individually, N-Ter 2 and N-Ter 3 inhibited migration by approximately 20% (Figure 5B). When combined, these peptides produced an additive effect, resulting in ∼40% inhibition comparable to that observed with the full N-Ter 1 peptide.

The C-terminal domain of STIM1 also plays a role in trans-endothelial migration. Targeting C-Ter^out^ STIM1_PM_ with a specific antibody (anti-STIM1 C-Ter 1) significantly inhibited JOK-1 migration by approximately 40%, an effect mirrored by the C-Ter 1 peptide (Figure 5C). Notably, an additive inhibitory effect was observed when combining antibodies targeting both the N- and C-terminal domains of STIM1 (Figure 5B). We hypothesize that the effects observed with antibodies targeting the N- and C-terminal domains of STIM1 are due to a combination of inhibition of constitutive calcium entry and disruption of interactions between STIM1 protein pools at the membranes of B cells and endothelial cells.

These findings demonstrate, for the first time, that CXCL12-induced trans-endothelial migration of JOK-1 cells is dependent on STIM1_PM_, regardless of its membrane orientation.

### Cell Specific Localization of STIM1_PM_ Influences Trans-Endothelial Migration of JOK-1 Cells

In all previous experimental conditions, whether assessing JOK-1 migration through a transwell filter or across an endothelial monolayer, only the lymphocytes were pre-incubated with STIM1-targeting peptides or antibodies. As previously reported (Figure 2), STIM1_PM_ N-Ter_out_ is expressed in both cell line types whereas STIM1_PM_ C-Ter_out_ is only present on Jok cell line. To determine the respective contributions of STIM1_PM_ in each cell type, we evaluated the effects of separate or simultaneous incubation of endothelial cells and JOK-1 cells with STIM1-targeting agents.

Incubation of endothelial cells with the anti-STIM1 N-Ter 1 antibody (10 µg/mL) inhibited JOK-1 trans-endothelial migration by 20% (Figure 6A), a level of inhibition comparable to that observed when only JOK-1 cells were treated. Interestingly, simultaneous treatment of both JOK-1 and endothelial cells with the antibody resulted in an additive effect, with approximately 45% inhibition of migration (Figure 6A).

**Figure 6.**
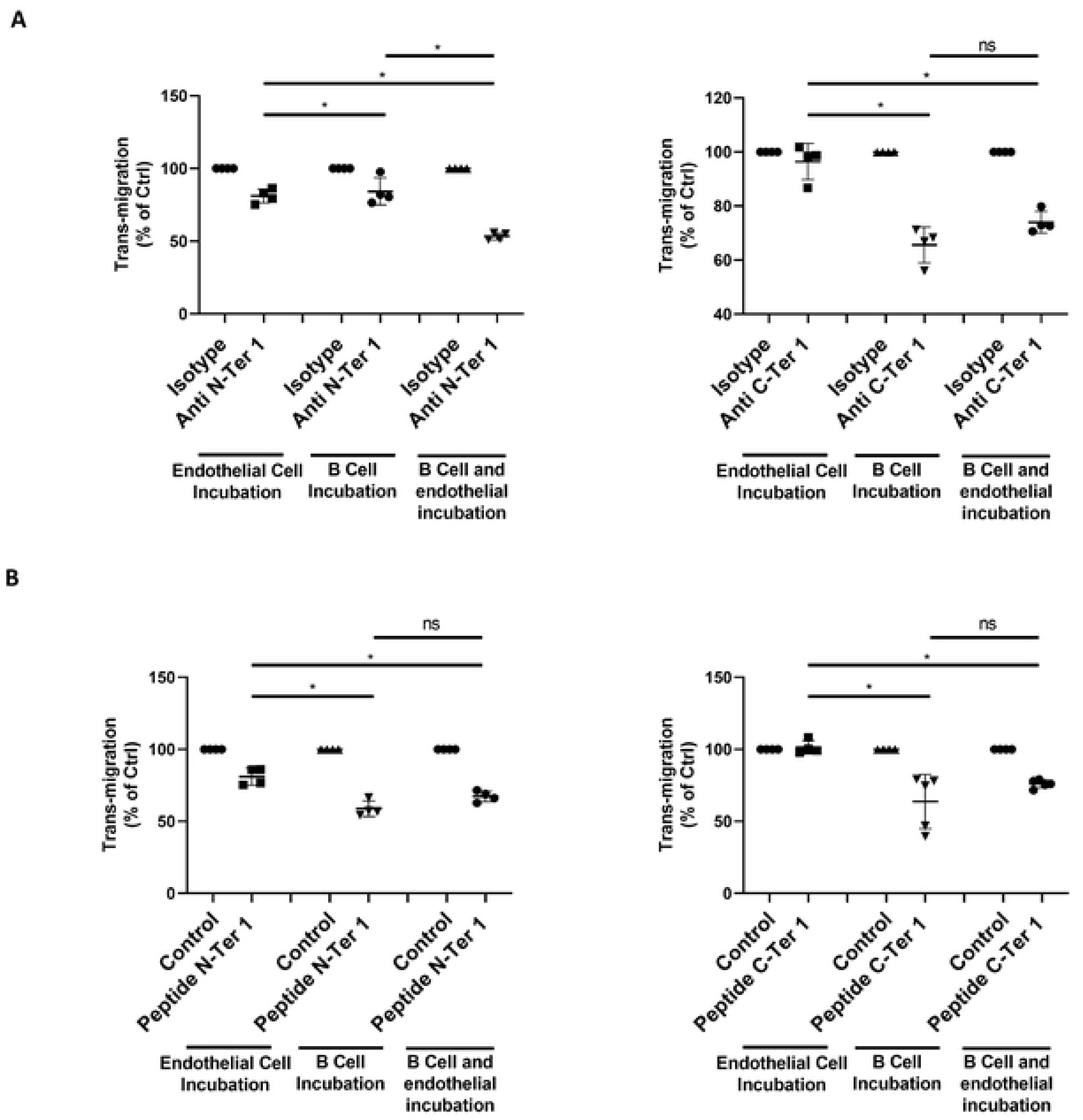
Trans-Endothelial Migration Is Differentially Modulated by STIM1_PM_ Depending on Its Localization in B Cells or Endothelial Cells. **A–B.** Effect of anti-STIM1 antibodies (**Panel A**: N-Ter 1, clone GOK, BD; C-Ter 1, clone CDN3H4, Santa Cruz; both at 10 µg/mL, *N*=4) and STIM1 peptides (**Panel B**: N-Ter 1, aa 63–200; C-Ter 1, aa 637–672; both at 10 µg/mL, *N*=4) on CXCL12-induced trans-endothelial migration of JOK-1 cells. In each experiment, JOK-1 and HUVEC cells were incubated either separately or simultaneously with antibodies or peptides for 1 hour. Statistical analysis: non-parametric *t*-test. *P*<0.05 (*), P<0.01 (**), P*<0.001 (\*\*\**).

When using the STIM1 N-Ter 1 peptide, migration was inhibited whether endothelial cells or JOK-1 cells were treated. However, no additive effect was observed when both cell types were treated simultaneously. In contrast, no inhibitory effect was observed as expected when only endothelial cells were treated with the anti-STIM1 C-Ter 1 antibody. Moreover, no significant difference in migration inhibition was observed between JOK-1-only and dual-cell-type treatments with this antibody (Figure 6B). Furthermore, no significant inhibition was observed when endothelial cells were treated with the C-Ter 1 peptide alone, and no additive effect was detected when both endothelial and B cells were treated (Figure 6B).

## Discussion

In this study, we confirmed the presence of STIM1_PM_ at the plasma membrane of both B lymphocytes and endothelial cells and investigated its role in B cell migration using an *in vitro* model of CXCL12-induced chemotaxis. For the first time, we demonstrated that STIM1_PM_ exists in B cells in two distinct topologies N-Ter_out_ (type I) and C-Ter_out_ (type II) and that this dual orientation contributes to the regulation of constitutive calcium entry and migration of B cells depending on its localization in B cells or endothelial cells.

Consistent with previous findings in human B cells (Debant et al., 2019) and in Panc1-WT (Buscaglia et al., 2026), we confirmed that STIM1_PM_ regulates constitutive calcium entry (CCE) but does not participate in the regulation of SOCE. These results were reproduced in the B cell line used in this study, further supporting the role of STIM1_PM_ in CCE regulation. Our findings support a role for STIM1_PM_ in chemokine-induced B cell migration, particularly in response to CXCL12. This regulation does not appear to involve modulation of the CXCL12-induced calcium signal, but rather is likely mediated through its involvement in CCE and the potentially through regulation of cell–cell interactions during transmigration.

### STIM1_PM_ : A Dual-Topology Protein Regulating Constitutive Calcium Entry in B Cells

This study is the first to describe STIM1 as a dual-topology protein at the plasma membrane of B cells, adopting either a type I orientation (N-terminal extracellular) or a type II orientation (C-terminal extracellular). Previous studies investigating STIM1_PM_ have consistently assumed a type I topology for this fraction of STIM1 (Debant et al., 2019; Mignen et al., 2007; Williams et al., 2001). However, recent work from our group has demonstrated that STIM1_PM_ is inserted into the plasma membrane in both orientations and can form antiparallel dimers (Buscaglia et al., 2026). In this context, both topologies of STIM1_PM_ were shown to regulate CCE. Additionally, glycosylation mutants revealed that STIM1_PM_ membrane localization is glycosylation dependent.

In the present study, flow cytometry using domain-specific antibodies confirmed the presence of dual-topology STIM1_PM_ in B cells. However, this dual topology was not observed in all B cell lines (e.g., RAMOS) or in HUVEC endothelial cells. As early as 2007, STIM1_PM_ was implicated in regulating ARC channels composed of Orai1 and Orai3, activated by arachidonic acid (Mignen et al., 2007; Thompson and Shuttleworth, 2012). More recently, STIM1_PM_ was shown to regulate CCE, involving Orai1 and TRPC1 channels, and this pathway was found to be upregulated in pathological B cells from CLL patients (Debant et al., 2019). Here, we demonstrate for the first time that STIM1_PM_ regulates CCE in B cells regardless of its membrane orientation, confirming observations made in pancreatic cancer cell lines (Buscaglia et al., 2026).

Furthermore, neither topology of STIM1_PM_ was involved in SOCE regulation, consistent with findings in CLL B cells (Debant et al., 2019) and pancreatic cancer cells. While STIM1_PM_ in B cells adopts both type I and type II topologies and regulates CCE, it is present in endothelial cells exclusively in a type I orientation. Considering this dual topology, the subsequent part of our study explored the role of STIM1_PM_ in B cell migration in response to chemokines.

### STIM1_PM_ Regulates Chemokine-Dependent B Cell Migration Without Modulating the Associated Calcium Signal

To date, very few studies have investigated the role of STIM1 in B cell migration in response to chemokine stimulation, particularly CXCL12. CXCL12 binds to its receptor CXCR4, triggering various intracellular signaling pathways involved in chemotaxis, including calcium signaling in multiple cell types.

In this study, we confirmed that CXCL12 induces a SOCE-type calcium influx in B cell lines, consistent with the known involvement of STIM1_ER_ in this process (Latour et al., 2018; Zuccolo et al., 2018). However, as previously reported (Debant et al., 2019), our findings demonstrate that STIM1_PM_ does not regulate SOCE and is not involved in the calcium signal triggered by CXCR4 activation. STIM1_PM_ does not influence the CXCL12-induced calcium signal, its role in B cell migration may be calcium-independent or at least SOCE independent. Blocking STIM1_PM_ N-Ter or C-Ter extracellular domains on B cells or blocking N-Ter domains on endothelial cells significantly inhibited CXCL12-induced migration. These results strongly support the involvement of STIM1_PM_ in chemokine-driven B cell migration and trans-endothelial migration. The effects of N-terminal peptides on migration mirrored those of the antibodies. However, an additive effect was observed when combining N-Ter 2 and N-Ter 3 peptides (both are fragments of N-Ter 1) suggesting that modulation of the EF and SAM domains may have distinct impacts on migration. Further experiments using antibodies targeting these fragments are needed to confirm this hypothesis. Nothing is known about the function of these STIM1 domains when exposred to the extracellular space for STIM1_PM_.

While both molecules inhibited B cell migration, the underlying mechanism remains therefore unclear. Taken together, these findings suggest that the N-terminal domain of STIM1, particularly the EF/SAM regions, plays a key role in regulating B cell migration. The C-terminal domain may also contribute, although no specific functional domain has yet been identified.

### STIM1_PM_ May Regulate B Cell Migration Through ECC and Cell–Cell Interactions

Based on the collective findings of this study, we propose several hypotheses to explain the role of STIM1_PM_ in B cell migration, which will require further validation.

First, we hypothesize that CCE may contribute to the basal activation of calcium-dependent signaling pathways involved in CXCL12/CXCR4-mediated B cell migration, such as the RhoA, PI3K, and MAPK pathways (Chatterjee et al., 2014; Latour et al., 2018). In a previous study, we showed that blocking CCE via an antibody targeting the N-terminal domain of STIM1 inhibited constitutive ERK phosphorylation in B cells from CLL patients (Debant et al., 2019). However, in the present study, the effects of some N-Ter antibodies and peptides on CCE were not similar but all have the same inhibitory effects on migration. This result making it difficult to attribute the observed migration effects solely to CCE modulation. Nevertheless, we cannot exclude a role for CCE in B cell migration, especially since CCE is also regulated by C-Ter_out_ STIM1_PM_.

Second, we propose that STIM1_PM_ may regulate cell/cell interactions during B cell migration. STIM1_PM_ was initially identified as a type II plasma membrane protein involved in B cell/stromal cell interactions (Oritani and Kincade, 1996). It is therefore plausible that interactions between the N-Ter or C-Ter domains of STIM1 proteins on adjacent cells contribute to migration. Although nothing is known about N-Ter STIM1_PM_ interactions, *in vitro* studies have shown that EF/SAM domain peptides can form calcium-dependent dimers (Stathopulos et al., 2006, Qiu et al., 2025, Narayanasamy et al., 2024). Also, STIM1 C-Ter domain can interact among them in the presence of Ca^2+^ in the ER (Shrestha et al., 2022 Nebel et al., 2007) these interactions contribute to the activation mechanisms of STIM1 to prevent the activation of STIM1. We hypothesize that blocking these domains with antibodies or peptides may disrupt B cell/endothelial or B cell/B cell interactions when only considering N-Ter, thereby impairing trans-endothelial migration. Notably, migration inhibition was observed even when only B cells were treated, suggesting that B cells-B cells interactions may also play a role in this model. This hypothesis remains speculative, as no studies to date have examined B cells-B cells interactions during chemokine-induced migration in the *in vitro* transwell model. We also consider the possibility that the C-Ter domain of STIM1 contributes to cell–cell interactions. The C-Ter region of STIM1 is known to mediate interactions with itself and with Orai1 channels via the CAD/SOAR domain (Prakriya and Lewis, 2015 (Shrestha et al., 2022, Nebel et al., 2007).

In this study, targeting the C-Ter domain with antibodies or peptides inhibited B cell migration. However, given the absence of C-Ter^out^ STIM1_PM_ in endothelial cells, we do not favor the hypothesis of C-terminal-mediated B cell–endothelial cell interactions. Instead, we propose that C-terminal domain interactions may play a role specifically in B cell/B cell interactions.

### STIM1_PM_-Mediated Inhibition of B Cell Migration: Independent of ECC and Cell–Cell Interactions?

An intriguing hypothesis that warrants further investigation is that STIM1_PM_ may regulate B cell migration through calcium-independent signaling pathways and like mechanisms described for integrins. Integrins are membrane proteins involved in cell–cell interactions, including lymphocyte/endothelial adhesion, and are essential for lymphocyte adhesion during trans-endothelial migration (Laufer and Legler, 2018). Through their intracellular domains, integrins interact with actin filaments and regulate adhesion by modulating actin cytoskeleton dynamics and activating key migration regulators such as RAS and FAK (Vicente-Manzanares et al., 2009; Palecek and Ubbels., 1997). It is therefore plausible that STIM1_PM_, like integrins, may participate in actin cytoskeleton reorganization. To explain the effects of targeting the N-Ter domain of STIM1_PM_ on migration, future studies should explore whether STIM1_PM_ modulates intracellular signaling pathways involved in migration (such as cytoskeletal remodeling) independently of CCE. Binding of antibodies or peptides to the N-terminal domain may induce conformational changes that propagate to the C-Ter domain, as described during SOCE activation (Vig et al., 2006), potentially altering downstream signaling.

An additive inhibitory effect was observed when B cells were treated with both N- and C-Ter STIM1-targeting antibodies, suggesting that these domains may regulate migration through distinct mechanisms. The N-terminal domain may influence cell/cell interactions and calcium signaling, while the C-terminal domain may regulate CCE and calcium-dependent signaling pathways involved in migration. Future studies should aim to dissect the specific roles of each STIM1_PM_ domain and the cellular processes they influence. This will require the development of molecular tools capable of distinguishing between STIM1_PM_-mediated effects on cell/cell interactions and those on CCE.

Taken together, our findings strongly support a role for STIM1_PM_ in CXCL12-induced migration of JOK-1 B cells. We propose that CCE contributes to chemokine-driven migration. In disease context, targeting STIM1_PM_ with antibodies or peptides may prevent B cell migration.

## CRediT authorship contribution statement

Nelig Le Goux Conceptualization, Methodology, Investigation, Data curation, Formal analysis, Writing - original draft, Writing - review & editing. Paul Buscaglia Data curation, Anthony Mainguy Data curation, Maella Hus Data curation, Mathieu Blery Conceptualization, Supervision, Olivier Mignen Conceptualization, Funding acquisition, Supervision, Investigation, Methodology, Validation, Writing - review & editing.

## Declaration of Competing Interest

The authors declare that they have no known competing financial interests or personal relationships that could have appeared to influence the work reported in this paper.

## Data availability

Data will be made available on request.

## Acknowledgements

We acknowledge theCalciscreen platforms, as well as the start-up Kalsiom SAS, for access to their facilities. Author NLG is supported by the “Allocations de Recherche Doctorale” (Région Bretagne).

## Materials and Methods

### Cell Culture

JOK-1 cells (ATCC) were cultured in RPMI 1640 medium (Sigma-Aldrich) supplemented with 10% fetal bovine serum (FBS) and 0.5% penicillin/streptomycin at 37°C in a 5% CO₂ atmosphere. HUVECs were cultured under the same conditions using Endothelial Cell Growth Medium 2 Kit (PromoCell, C-22111) supplemented with 0.5% penicillin/streptomycin.

### *In Vitro* Cell Migration Assays

#### Transwell Migration

Migration assays were performed using 6.5 mm diameter transwell inserts with 5 µm pore size (Corning). JOK-1 cells (2.5 × 10⁵) were seeded in the upper chamber and incubated for 24 h at 37°C, 5% CO₂ in RPMI 1640 medium (BioWest, with L-glutamine and 25 mM HEPES, ref L0495-500) supplemented with 1% penicillin/streptomycin and 0.5% BSA. Cells were pre-incubated for 1 h with one of the following treatments:

- Mouse monoclonal anti-STIM1 antibodies: C-Ter 1 (clone CDN3H4, Santa Cruz, sc-66173), N-Ter 1 (clone GOK, BD, 624094), N-Ter 2 (clone BY-12, Diaclone), N-Ter 3 (clone BM-4, Diaclone)
- STIM1 peptides: N-Ter 1 (EF-SAM, aa 63–200), N-Ter 2 (1A, aa 132–160), N-Ter 3 (2B, aa 96–128), C-Ter 1 (C2, aa 637–672)

The lower chamber contained 600 µL of RPMI 1640 with 1% penicillin/streptomycin, 0.5% BSA, and 200 ng/mL CXCL12 (Recombinant Human/Rhesus Macaque/Feline CXCL12/SDF-1α, Bio-Techne, 350-NS-010). Migrated cells in the lower chamber were quantified by flow cytometry (Navios 1, Beckman Coulter). Migration percentages were normalized to isotype or vehicle controls.

#### Trans-Endothelial Migration

The same transwell setup was used. HUVECs (1 × 10⁵) were seeded in the upper chamber and cultured for 24 h. JOK-1 or HUVEC cells were pre-incubated for 1 h with the treatments listed above. The lower chamber contained the same medium and CXCL12 concentration as described above. Migrated cells were quantified as previously described.

### Calcium Influx Measurement Following CXCL12 Stimulation

Cells were loaded with 2 µM Fura-2/AM and 2 µM pluronic acid for 60 min at 37°C in buffer A (135 mM NaCl, 1.8 mM KCl, 1 mM MgCl₂, 10 mM HEPES, 10 mM glucose, pH 7.4) supplemented with 5 mM CaCl₂. Fura-2 was excited alternately at 340 and 380 nm using a polychromator (TILL Photonics), and fluorescence emission was recorded at 510 nm using an IX71 fluorescence microscope (Olympus) equipped with a 415DCLP dichroic mirror and a 14-bit CCD camera (ExiBlue, QImaging). After baseline stabilization, cells were stimulated with 200 ng/mL CXCL12. The F340/F380 ratio was calculated using Metafluor 6.3 software (Universal Imaging), and calcium influx amplitude was expressed as ΔF/F₀.

### Measurement of Constitutive Calcium Entry (CCE) via Mn²⁺ Quenching

96-well plates were coated with Cell-Tak (Corning), and 5 × 10⁵ JOK-1 cells were adhered for 1 h at 37°C, 5% CO₂. Cells were loaded with Fura-2 QBT™ (Molecular Probes) according to the manufacturer’s instructions. Treatments (peptides, antibodies, inhibitors, or controls) were applied during the adhesion period. Fura-2 was then replaced with calcium-free buffer (135 mM NaCl, 5 mM KCl, 1 mM MgCl₂, 1 mM EGTA, 10 mM HEPES, 10 mM glucose, pH 7.4). Intracellular calcium changes were measured using a FlexStation 3™ (Molecular Devices, UK). Mn²⁺ quenching was monitored by excitation at 360 nm and emission at 510 nm.

### Measurement of SOCE

The same protocol as for CCE was used. After Fura-2 loading and treatment incubation, cells were washed and incubated in calcium-free buffer. Ratiometric calcium measurements (excitation at 340/380 nm, emission at 510 nm) were recorded using a FlexStation 3™. SOCE was induced by ER calcium store depletion using 2 µM thapsigargin (in buffer A), followed by the addition of extracellular calcium to a final concentration of 1.8 mM. SOCE amplitude and kinetics were analyzed for each condition.

### Flow Cytometry

For STIM1 surface expression analysis, 5 × 10⁶ cells were used per condition. Adherent cells were detached using enzyme-free dissociation solution (Thermo Fisher) for 5 minutes at 37°C. Cells were centrifuged at 300 × *g* for 5 minutes and resuspended in 50 µL PBS containing one of the following PE-conjugated anti-STIM1 monoclonal antibodies:

- N-terminal domain: N-Ter 1 (GOK, BD Biosciences, 2.5 µL, 10 µg/mL), N-Ter 2 (BY-12, Diaclone, 2 µL, 10 µg/mL), N-Ter 3 (BM-4, Diaclone, 2 µL, 10 µg/mL)
- C-terminal domain: C-Ter 1 (CDN3H4, Santa Cruz, sc-66173, 5 µL, 10 µg/mL) Isotype controls (0.5 µL) were used in parallel. Cells were incubated on ice for 20 minutes, washed three times with PBS, and analyzed using a Navios flow cytometer (Beckman Coulter Life Sciences). Median fluorescence intensity (MFI) was calculated using Kaluza 2.1 software (Beckman Coulter Life Sciences).

For CXCR4 and CXCR5 surface expression, the same protocol was followed. Cells were incubated for 20 minutes with anti-CXCR4 (1 µL, APC-conjugated, clone 12G5, BioLegend) and anti-CXCR5 (1 µL, BV421-conjugated, clone J252D4, BioLegend) antibodies.

## Notes

### Competing Interest Statement

The authors have declared no competing interest.

